# Recurrent connectivity underlies lateralized temporal processing differences in Auditory Cortex

**DOI:** 10.1101/2021.04.14.439872

**Authors:** Demetrios Neophytou, Diego Arribas, Robert B. Levy, Memming Park, Hysell V. Oviedo

**Author notes:** Corresponding author **Address correspondence to:** Hysell V. Oviedo, Department of Biology, The City College of New York, 160 Convent avenue, Marshak room 526, New York, NY 10031. Co-first authors (Demetrios Neophytou, Diego Arribas).

## Abstract

Brain asymmetry in the sensitivity to spectrotemporal modulation is an established functional feature that underlies the perception of speech and music. Recent studies in humans suggest separable neural mechanisms must underlie these cognitively complex functions. The left Auditory Cortex (ACx) is believed to specialize in processing fast temporal components of speech sounds, and the right ACx slower components. However, the circuit features and neural computations behind these lateralized spectrotemporal processes are poorly understood. To answer these mechanistic questions we use mice, an animal model that captures some relevant features of human communication systems. In this study, we screened for circuit features that could subserve temporal integration differences between the left and right ACx. We mapped excitatory input to principal neurons in all cortical layers and found significantly stronger recurrent connections in the superficial layers of the right ACx compared to the left. We hypothesized that the underlying recurrent neural dynamics would exhibit differential characteristic timescales corresponding to their hemispheric specialization. To investigate, we recorded spike trains from awake mice and estimated the network time constants using a quasi-Bayesian method to combine evidence from multiple weak signal-to-noise ratio neurons. We found longer temporal integration windows in the right ACx compared to the left as predicted by stronger recurrent excitation. Our study shows direct evidence that stronger recurrent synaptic connections lead to longer network time scales. These findings support speech processing theories that purport asymmetric integration time constants is a crucial feature of lateralization in auditory processing.

## Introduction

Social communication calls have myriads of constituent sounds that are temporally and spectrally dynamic. The auditory system must have processes in place to quickly decode and encode ethologically relevant features of an auditory signal to elicit an appropriate response. Asymmetry in sound processing (i.e. lateralization) has long been proposed to be critical in the dynamic processing of speech sounds. Human studies have shown that the left superior temporal gyrus (STG) is more capable of integrating information over a shorter timescale and plays a greater role in speech perception and phonological processing than the right (1-3). On the other hand, the right STG has longer integration windows to potentially subserve the processing of suprasegmental information (4, 5). The unanswered question remains: What neural mechanisms underlie these differences in temporal integration?

Studies of animal models can provide more mechanistic insight regarding the function and motif of auditory circuits. Circuit mapping of the mouse ACx has shown that the synaptic organization of Layer 3 (L3) differs between the two hemispheres. In the left ACx, principal neurons in L3 receive out-of-column excitatory input from L6 cells located in higher frequency bands, whereas in the right ACx the same pathway has balanced frequency projections across the tonotopic axis. This lateralized synaptic organization is in turn associated with differences in sound-evoked activity in L3 (6). Behavioral studies in gerbils have reported asymmetries in temporal processing. Lesions in the right ACx impact discrimination of frequency sweep direction, suggesting it plays a role in processing global temporal cues. Whereas lesions in the left ACx impact discrimination of gap durations, implying a role in processing local temporal cues (7). Here, we test directly whether there are differences in the temporal integration properties between the Auditory Cortices and dissect the underlying circuit dynamics. To assess cortical circuit mechanisms that could underlie hemispheric asymmetry in temporal processing, we used circuit-mapping techniques to screen for connectivity differences in excitatory pathways. We show surprising differences in recurrent connectivity between the hemispheres, particularly in the superficial layers. To examine how these recurrent asymmetries translate into differences in temporal integration we recorded spontaneous spikes from awake mice, and developed a quasi-Bayesian method to estimate time constants from spike trains. We use the dichotomized Gaussian model to reproduce autocorrelations and generate surrogate spike trains that were used to estimate uncertainty and provide a belief distribution over the network time constant. Applying this method to data recorded from superficial layers of the left and right ACx, we found significant differences in temporal integration consistent with the timescale of hypothesized lateralized auditory signal processing. Numerous models of neural architectures have been proposed to account for observed differences in integration timescales throughout the brain. Here we show for the first time direct evidence of differences in synaptic circuit organization that translate into distinct temporal integration windows.

## Results

### Lateralized connectivity motifs are found in every layer of the Auditory Cortices

To screen for hemispheric differences in the organization of excitatory pathways, we used glutamate uncaging based Laser Scanning Photostimulation (LSPS; (8)). We performed voltage-clamp recordings on principal neurons in Layers 2-6 of the left (n = 237) and right (n = 239) ACx, (Figure 1A-B). The uncaging stimulus grid covered the entire primary ACx and all cortical layers (total area of 1.125*1.125mm, 256 stimulation sites/map pixels). We specifically focused on measuring intracortical sources of excitatory synaptic input by holding the membrane potential at the reversal for inhibition (−70mV). To capture a global view of potential synaptic connectivity differences in each cortical layer, we performed a statistical comparison of the population data underlying corresponding map pixels from each hemisphere. We found statistically significant differences in the strength and organization of synaptic input between most layers of the auditory cortices (Figure 1C). To assess the likelihood that these differences are by chance, we randomly assigned each pixel to the left or right ACx. We largely found fewer significant differences between the hemispheres when the pixels were randomized (Figure 1D).

**Fig 1.**
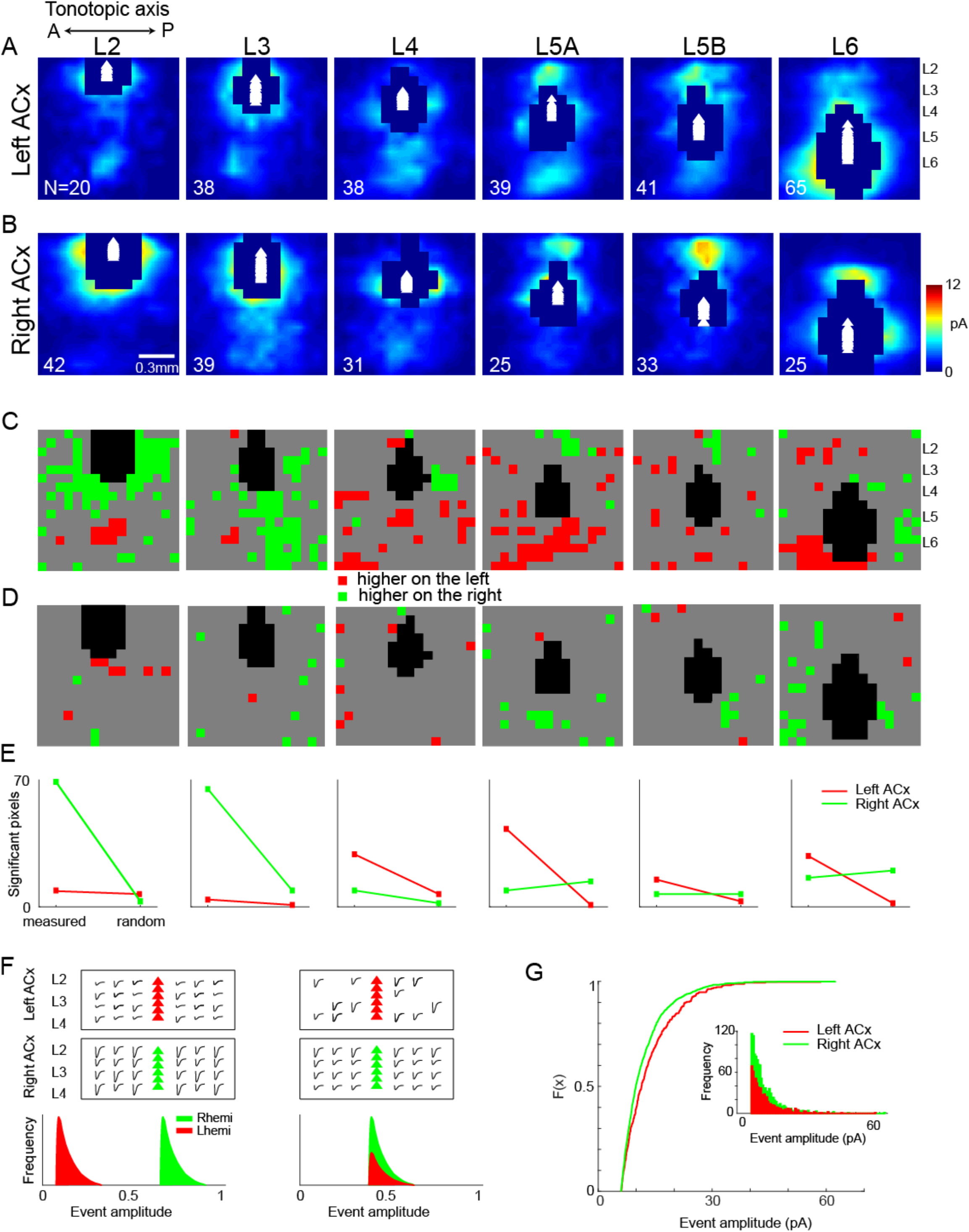
Summary of excitatory pathways in the left and right ACx across all cortical layers. (A, B) Patterns of excitatory synaptic input were recorded using LSPS as described in the text. Maps were averaged over all cells in each layer and then interpolated for clarity. Masked areas indicate direct hits in >50% of cells for that region. White triangles denote location of somata. Laminar boundaries were defined with respect to the fractional distance from the L1/L2 boundary to the white matter (WM). L5A and L5B were defined as the upper and lower 50% of L5, respectively. N indicates number of recorded cells for each panel. (C) Pixel-wise significance maps (p<0.05, unpaired 2-tailed t-tests) for response amplitude in left vs. right ACx. *Red* and *green* denote significantly higher average response in the left and right ACx, respectively. *Gray* denotes no significant difference. (D) Same as C but maps from both hemispheres were pooled and assigned to two groups at random with the same total n shown in A,B. (E) Graphs of the significant pixel counts for measured (C) and random (D) comparisons. (F) Models depicting potential synaptic mechanisms underlying the right ACx’s higher excitatory connectivity in superficial layers compared to the left ACx. The left panel shows both cortices with a similar pool of presynaptic sources of EPSCs (traces) projecting onto postsynaptic targets (triangles), but the distribution of event amplitudes differs. The right panel shows the left and right ACx differ in their pool of presynaptic inputs, but have a similar event amplitude distribution. (G) Observed empirical cumulative distribution of event amplitudes in superficial layers of the left and right ACx. Inset shows frequency distribution.

A large source of presynaptic input to L2 in the right ACx arose intralaminarly and from other superficial layers (Figure 1A-C). In contrast, in the left ACx the most influential presynaptic pathway to L2 arose from deep layers (L5/6), as was reported previously (9). There are also hemispheric differences in the organization of input to L3, which were also previously described in detail (6). Differences in projections to L4 were more broadly distributed between the left and right ACx (Figure 1A-C). Layers 5A and B are functionally distinct in the ACx, and the organization of their intracortical synaptic input appears to be lateralized. In the left ACx there was greater bottom-up input to L5A compared to the right ACx. Conversely, there was greater top-down input to L5B in the right ACx compared to the left ACx (Figure 1A-C). Finally, L6 had complementary patterns of intralaminar synaptic input along the tonotopic axis: the left ACx had greater input arising from higher frequency bands, and the right ACx from lower frequency bands (Figure 1A-C). A comparison of the number of measured and random pixels for each layer revealed that the most abundant differences in synaptic input between the hemispheres arose in the superficial layers of the left and right ACx (Figure 1E). Several synaptic mechanisms could underlie the observation of more significant hotspots in superficial layers in the right ACx compared to the left. One possibility is that each hemisphere has a similar pool of presynaptic inputs in these layers, but individual synaptic events are larger in the right ACx compared to the left (Figure 1F left). Another possibility is that the distribution of the synaptic events’ amplitudes is similar between the hemispheres, but there is a larger pool of presynaptic inputs in the right ACx compared to the left (Figure 1F right). To disentangle these possibilities we analyzed the distribution of event amplitudes in superficial layers (L2-4). We randomly chose the same number of cells to analyze from each hemisphere (n=96), and the threshold for synaptic events was set to 4 standard deviations above the average baseline. Fewer synaptic events met the threshold criteria in the left ACx (688) compared to the right (1148). The observed cumulative distribution of synaptic event amplitudes shows that there is a significant difference between the hemispheres (Figure 1G; p=0.0017, kstat=0.1198, n=96, two-sample Kolmogorov-Smirnov test). This supports the prediction that there is a larger pool of presynaptic inputs projecting to postsynaptic targets in superficial layers of the right ACx.

### Lateralized recurrent connectivity in superficial layers of the Auditory Cortices

Neural circuits composed of short time constant units (i.e. neurons) can effectively have long temporal memory and computation by forming a long feedforward chain or a concise recurrent feedback loop (10). Therefore, we investigated whether the synaptic connectivity differences translate into systematic differences in the recurrent interlaminar feedback. To compare the relative strength of interlaminar pathways between the hemispheres we computed connectivity matrices for the left and right ACx. These input-output matrices (presynaptic-postsynaptic) summarize the organization of local excitatory networks succinctly (11, 12). We ordered maps according to the cortical depth of the soma, summed the two dimensional LSPS-derived input maps for each cell over the tonotopic axis (across the horizontal dimension) to produce vectors of input strength as a function of cortical depth (Figure 2A). Therefore, each neuron’s input vector represents presynaptic input from different laminar locations. Each row in the laminar connectivity matrix represents input to that specific laminar location, and each column represents synaptic output from that laminar location. Very local connections (<50microns) within each layer lie along the diagonal and were under-sampled due to direct excitation (11). In a previous study we determined that neuronal density and photoexcitability of individual cells did not significantly differ between the hemispheres; therefore we did not normalize maps by these factors (6).

**Fig 2.**
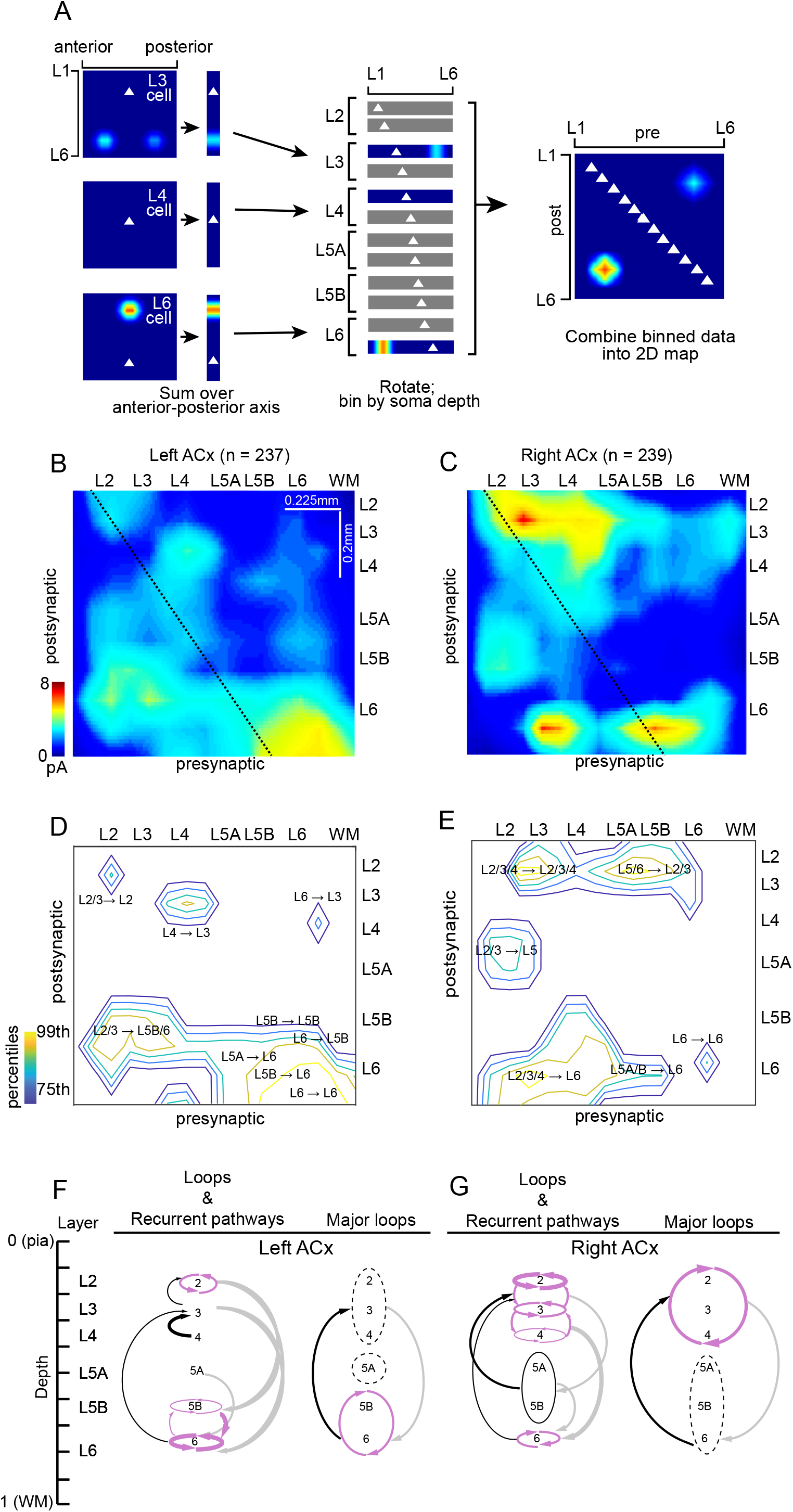
Recurrent connections are significantly stronger in superficial layers of the Right ACx. (A) Schematic diagram of laminar input-output maps (12). *Left*, two dimensional LSPS-derived input maps for each cell were summed over the anterior posterior axis to produce one dimensional maps of input strength vs cortical depth (vertical strips). Note that anteroposterior information is discarded, such that the two L6 input hotspots for the L3 cell (top) collapse to a single spot. Triangles denote soma location. *Middle*, the 1D maps were rotated 90 degrees (for graphic clarity), sorted by cortical depth of the soma, and binned (bin size = 80 μm). *Right*, the binned maps were combined into a single 2D map of presynaptic input location (x axis) vs. binned postsynaptic soma location (y axis). Maps were interpolated for display. (B) Input-output map for the left ACx, constructed as shown in A. Diagonal line indicates x=y with respect to cortical depth. The diagonal does not span the full x axis because recorded cell bodies (y axis) were confined between L2 and L6, whereas the stimulation grid (x axis) extended more broadly from L1 into the WM. (C) Same as B but for the right ACx. (D) Same map as in B but showing only pathways in the 75^th^ percentile and above in the left ACx. (E) Same as D but for the right ACx. (F) Summary of loops and pathways in the 75^th^ percentile and above in the left ACx, and right ACx (G). In F and G the arrow thickness indicates strength of the pathway, ascending pathways are shown in black, descending in gray, recurrent in violet, and open loop in dashed line.

The most striking asymmetry in the laminar connectivity matrices was the stronger synaptic connections in superficial layers of the right ACx compared to the left (Figure 2B-C). The fractional input and output of layers 2-4 was significantly greater in the right ACx (p = 0.0079 for input, p = 0.0411 for output). In the deeper layers, input to L6 was significantly greater in the left ACx (p<<0.001), but there was no hemispheric difference in the output. Significant hemispheric differences in intralaminar and interlaminar loops (input arising and returning to the same layer(s)) were a major theme in the organization of auditory circuits. Using the input/output laminar connectivity matrices of the left and right ACx, we examined pathways in the 75^th^ percentile to capture the most significant trends. In the left ACx we observed strong intralaminar recurrent connectivity only in L6 and to a lesser extent in layers 2 and 5B (Figure 2D). The strongest interlaminar recurrent connections were observed in deep layers, where L5B and 6 form nested loops. These loops receive input from almost all layers (except L4), and its output is relayed to L3 (Figure 2F). In contrast to the sparse prevalence of recurrent connections and loops in the left ACx, these were the dominant motifs in the right ACx. All cortical layers (except L5) in the right ACx are part of nested loops: at the lowest level they have intralaminar recurrent connections, followed by recurrent connectivity to neighboring layers, and at the highest level long-range recurrent connections that couple superficial and deep layers (Figure 2E, G). Taken together, these widespread loops of recurrent connections suggest the right ACx may have different temporal filtering properties and enhanced sensory memory capabilities induced by network structure compared to the left ACx.

### Network time constant is longer in superficial layers of the right Auditory Cortex

To quantify the difference in the network dynamics impacted by differential recurrent connectivity, we measured the time constant of the spontaneous neural activity in awake animals. Time constant provides a first order approximation of the dynamics by measuring how quickly the correlation in the neural activity decays over time. Without any recurrent connectivity, the time constant of single neuron activity is at the scale of membrane time constant and delays, while with recurrent connectivity, the fluctuations in the network activity can decay much slower (i.e., longer time constant) due to effective self-excitation. Therefore, we studied whether hemispheric differences in recurrent connectivity translate into a difference in the network time constants reflected in the single neurons’ spontaneous activity in the absence of auditory stimuli. We performed cell-attached recordings in superficial layers (150-450 microns below the cortical surface) of the left and right ACx in head-fixed awake mice because anesthesia can reduce the contribution of network activity (13). Neurons in both auditory cortices were active in the absence of auditory stimuli and displayed activity that suggested the presence of temporal correlations (Fig. 3C).

**Fig 3.**
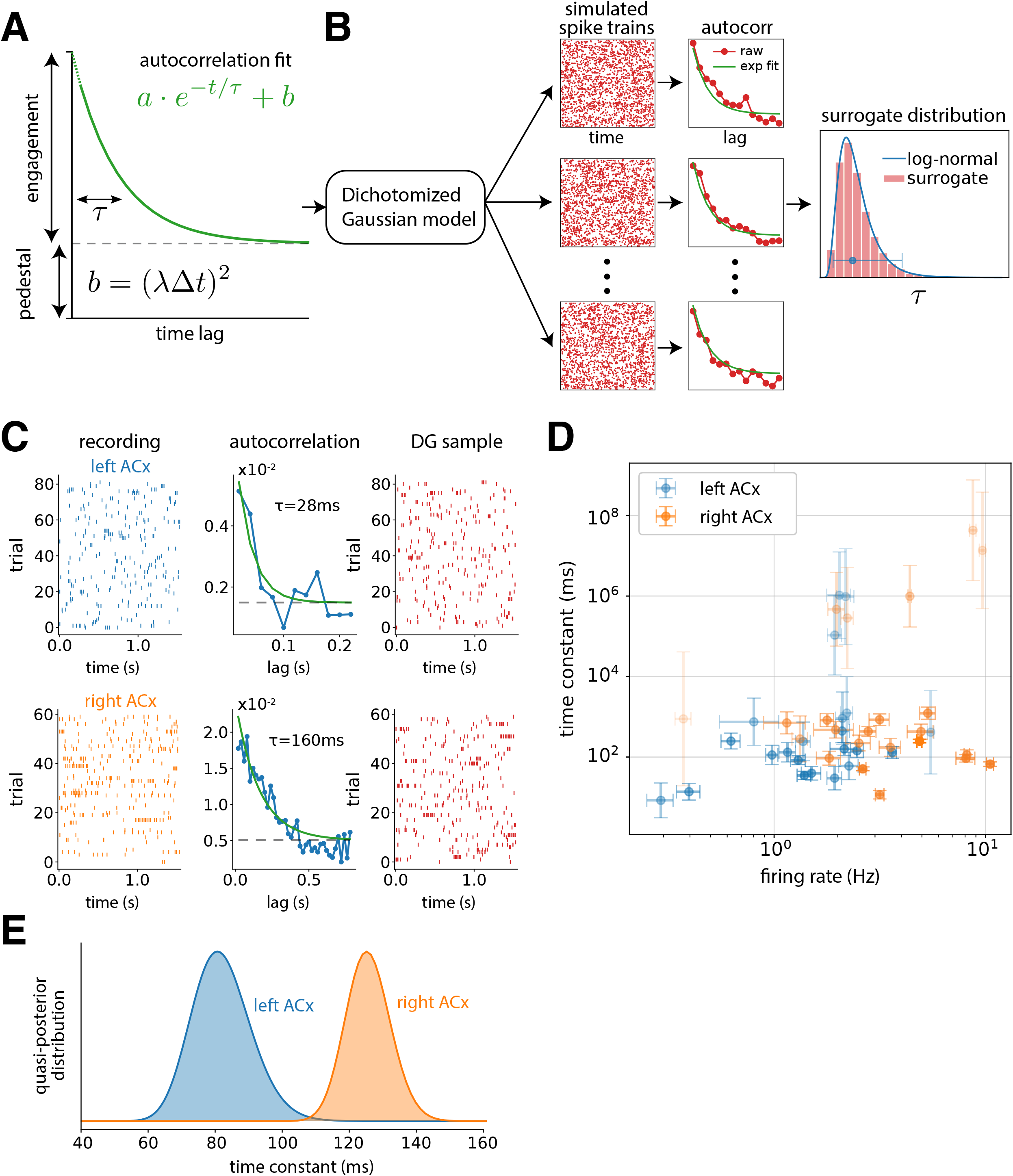
Time scales of neural activity are longer in the right ACx. (A) Schematic of the fitted exponential decay and its parameters. (B) Sketch of the procedure to estimate the bias and uncertainty in the time constant of single neurons. We use the DG model with exponential autocorrelations to generate surrogate data, re-fit exponentials and extract surrogate time constants. Finally, we fit a lognormal distribution to the surrogate time constants to estimate the bias and variance. (C) Example left and right ACx data, autocorrelations and sample of surrogate data generated from DG model with the extracted autocorrelation. (D) Bias corrected time constant with its uncertainty against mean firing rate for each neuron. Time constant error bars are the standard deviation of the surrogate lognormal distribution. Firing rate error bars are the standard error over trials. (E) Time constant posterior distributions obtained by integrating the single neuron observations for each hemisphere.

We extracted the time constant of individual cells by fitting an exponential decay with a pedestal to their autocorrelograms (Fig. 3A). However, precise quantification of time constants from spike trains and their autocorrelograms can be challenging (14, 15) especially for neurons with low firing rates or low degrees of engagement with the network. In such low signal-to-noise-ratio (SNR) regime where spike trains (signal + noise) are not very informative about the network time constant (signal), the estimated time constant can exhibit large bias and variance. Unfortunately, bootstrapping or resampling procedures cannot practically correct for these issues. For example, if there are only a handful of spikes, bootstrap samples may consistently provide very small time constants such that the variance of the estimator is close to zero. However, in this low firing rate regime, the uncertainty of the estimator should be large if we properly accounted for the spiking noise – resampling low SNR data combined with a strongly biased estimator can create an illusion of a (misleading) high SNR conclusion.

To better estimate the bias and variance of the time constant estimator itself, we used a method for generating spike trains from a parametric surrogate distribution (Fig. 3B, Methods). Briefly, for each neuron, we use the exponential fit of the data’s autocorrelation to build a probabilistic model (see Methods) that we use to generate data that reproduces the autocorrelation. We generate surrogate data from this model matching the length and sampling rate of the recording. By construction, the generated data reproduces the firing rate and autocorrelation of the original data on average, while single realizations will show variability determined by the assumed noise. We then re-fit exponential decays to the surrogate data and extract surrogate time constants from each of the realizations. Using this surrogate time constant distribution, we correct the bias by assuming that the bias is identical between data and surrogate fits, and obtain a time constant estimate for each neuron together with an uncertainty around this value (Fig. 3D) that captures the statistical variability of the estimator (e.g., represented by the log-variance of the estimator). The extracted time constants and their uncertainties spanned multiple orders of magnitude and there was no clear correlation with the average firing rates unlike the bootstrap estimation (data not shown).

To systematically aggregate the unequally informative neurons, we used a quasi-Bayesian method that accounts for the bias and uncertainty in the time constant estimation for each neuron (Fig. 3B, Methods). Neurons with high uncertainty (low SNR estimate) automatically provide relatively weak evidence about the network’s time constant. Hence, our method does not apply arbitrary criteria to discard neurons with weak autocorrelations, low firing rates or very long timescales. We then integrate the evidence from multiple neurons assuming a shared network time constant per hemisphere (Fig. 3E, Methods). The extracted network time constants were 82ms (66ms, 99ms) for the left ACx and 126ms (113ms, 138ms) for the right ACx (mean, (95% credible interval)). We found that the time constant of the right ACx network was significantly longer (Bayes factor=0.002. One posterior for all neurons compared to different posterior for each hemisphere). Our results suggest that differences in the recurrent connectivity in superficial layers of the Auditory Cortices translate into differences in time constants that might support lateralized computation of auditory stimuli.

## Discussion

Rodent studies of the last several decades have demonstrated that there are evolutionarily conserved mechanisms of vocalization processing. Behavioral and electrophysiological evidence supports the role of the left ACx in the perception of conspecific vocalizations (6, 16, 17) and detecting local features in auditory signals, whereas the right ACx specializes in global features (6, 7). Moreover, these processing asymmetries between the auditory cortices are associated with lateralized circuit-motifs (6). In this study, we show a novel neural mechanism whereby hemispheric differences in recurrent connectivity in superficial layers of the ACx translate into distinct temporal integration time windows. We showed that the synaptic organization, network dynamics, and function consistently exhibit lateralization. Using laser scanning photostimulation, we compared the organization and strength of excitatory pathways in the left and right ACx across all cortical layers. For each auditory cortex we combined the data across cells by building input-output maps. We found significantly stronger intralaminar and interlaminar connectivity between cells in the superficial layers in the right ACx than in the left, suggesting stronger recurrent loops (Figure 2). Stronger recurrent activity in the right ACx suggests a capacity for enhanced echoic memory: holding a brief memory of auditory signals. This feature would be necessary to extract auditory information at slower temporal timescales, such as prosody and intonation.

Recurrent connectivity and their positive feedback has long been conjectured to play a key role in persistent neural information processing by lengthening the temporal extent of information representation (10, 18, 19). Although there has been a bevy of theories and computational models (20-23) only indirect evidence exists correlating recurrent anatomical structure with observed persistent activity or longer time constants. Previous work has shown that timescales extracted from single neurons’ activity are related to cortical network organization in the primate visual processing hierarchy (24), and to the integration of signals in tasks such as working memory (25), reward guided choice (26, 27), and free choice (28). But there has not been a comparison of the naturally existing recurrent excitatory synaptic circuits and the corresponding network time constants within the same biological model system. In this work, we were able to exploit the lateralized recurrent circuit architecture of the ACx to make the desired direct comparison. The theory suggests that more excitatory recurrent feedback within the superficial layers of the right ACx should give rise to longer network time constant reflected in the neural activity. These differences in recurrent connectivity should impact the temporal structure of both evoked and spontaneous activities (29, 30); however, it is easier to interpret the neural dynamics in the absence of stimulus drive since stimulus dynamics influences the measured time constants. To test the hypothesis, we performed cell-attached recordings in superficial layers of the left and right ACx in awake mice and studied the autocorrelogram of the spontaneous neural activity. Consistent with the connectivity results, our statistical inference showed that the superficial layer network of the right ACx exhibits a ∼50% longer network time constant relative to the left. To the best of our knowledge, this is the first direct evidence that excitatory synaptic feedback predicts the spontaneous network dynamics that utilize the natural consistent variations provided by lateralization.

The consensus from decades of human language studies has been that speech decoding is carried out bilaterally with each hemisphere optimized for specific functions. Activity in the left STG has been consistently associated with processing of formant transitions, and perception of speech content is affected by degradation of temporal information; whereas activity in the right STG is associated with processing of intonation contours and is affected by degradation of spectral information (1, 31). However, the underlying neural mechanisms have remained controversial and difficult to prove in humans. One proposed mechanism that could subserve lateralized processing is neuronal ensembles in each hemisphere with different integration time constants: neurons in the left ACx could preferentially integrate information on shorter time scales, and neurons in the right ACx could preferentially integrate information on longer time scales (4). Our results support this simultaneous multiscale temporal analysis. Mouse vocalizations are composed of syllables (i.e. a sound unit separated from other sound units by silence (32), which themselves contain one or several pitch trajectories. The duration of individual pitch trajectories within a syllable emitted by adult male mice can range from 30-90ms (33, 34). This range of durations is in line with the distribution of integration time constants we observed in the left ACx (Fig. 3E). On the other hand, the right ACx’s time constants are consistent with intersyllabic intervals in mouse vocalizations (140ms on average; (35)). These hemispheric differences in processing timescales suggest that the left and right ACx are simultaneously processing segmental and supra-segmental information, respectively, and thereby extracting different acoustic cues from the same signal (4). While the results of our study support asymmetries in temporal integration, our previous work suggests that this is not the only neural mechanism underlying lateralized auditory processing. There are hemispheric differences in the tonotopic organization of auditory cortical circuits and responses to frequency sweeps, which suggest that there are also lateralized specializations for spectral processing (6). Overall, the emerging evidence is that there are neural specializations for division labor in both the temporal and spectral domain.

As the list of circuit asymmetries grows in the ACx it will be important to consider how they arise. Asymmetries in genetic programs that guide circuit assembly during development as well as hearing-experience are very likely sources of lateralized specializations. These forces can influence the asymmetric formation of both excitatory and inhibitory networks (36). The observation of greater recurrent connectivity in the right ACx potentially suggests a lower rate of pruning of synaptic contacts during the critical period compared to the left ACx. But it is unclear how layer-specificity would be achieved. In future studies we plan to dissect cellular and molecular mechanisms of enhanced recurrent connectivity in the right ACx. Although the results from this study are from an animal model, they support prevailing findings from the human literature, provide detailed neural mechanisms and therefore can potentially offer insight for investigations in humans (37).

## Methods

### Slice preparation and electrophysiology

We used CBA/J male mice aged 28-51. Animals were anesthetized and decapitated and the brains were transferred to a chilled cutting solution composed of (in mM): 110 choline chloride, 25 NaHCO_3_, 25 D-glucose, 11.6 sodium ascorbate, 7 MgCl_2_, 3.1 sodium pyruvate, 2.5 KCl, 1.25 NaH_2_PO_4_, and 0.5 CaCl_2_. We made horizontal slices to map synaptic connectivity along the anterior-posterior axis of the brain where tonotopy is represented in the ACx. We sliced using a 15-degree angle between the blade and the medial-lateral axis so that apical dendrites were parallel to the slice in the ACx. Slices were 300 μm thick and were transferred to artificial cerebrospinal fluid (ACSF) containing (in mM): 127 NaCl, 25 NaHCO_3_, 25 D-glucose, 2.5 KCl, 1 MgCl_2_, 2 CaCl_2_, and 1.25 NaH_2_PO_4_, aerated with 95% O_2_ 5% CO_2_. The slices were incubated at 34° for 20–30 minutes and then kept at room temperature during the experiments. Excitatory neurons located >50 μm below the surface of the slice were visualized using infrared gradient contrast optics and patched with electrodes (6–7 MOhm) containing the following intracellular solution (in mM) 128 K-methylsulfate, 4 MgCl, 10 HEPES, 1 EGTA, 4 NaATP, 0.4 NaGTP, 10 Na-phosphocreatine. The pH of the intracellular solution was adjusted to 7.25 and the osmolarity was 300 mOsm. Whole-cell recordings were made using a Multiclamp 700A amplifier (Axon Instruments, Molecular Devices, Sunnyvale, California, USA). Excitatory synaptic currents were measured at a holding potential of –70 mV, and for excitation profiles (see below) action potentials were recorded in the cell-attached configuration. We used the custom software package ephus ((38)http://www.ephus.org) for instrument control and acquisition written in Matlab (MathWorks, Natick, Massachusetts, USA).

### LSPS by glutamate uncaging

The ACSF for uncaging was supplemented with (in mM): 0.2 nitroindolinyl (NI)-caged glutamate (Tocris), 0.005 CPP (Tocris), and a final concentration of 4 CaCl_2_ and 4 MgCl_2_. For focal photolysis using UV flash to activate the caged glutamate compound, we used a 1 ms light stimulus consisting of 100 pulses from a pulsed UV laser (wavelength, 355 nm with a repetition rate of 100 kHz; DPSS Lasers, Santa Clara, California USA). The grid to stimulate the ACx had 16×16 uncaging spots with 75 μm spacing, which resulted in a mapping region of 1.125×1.125 mm. To avoid revisiting the vicinity of sites recently stimulated, we used a shifting-X pattern that was rotated and/or transposed between map iterations. Each stimulus trial contained a test pulse to measure electrophysiological parameters, and UV flashes were presented every 1 s. The stimulus grid was consistently aligned for each cell recorded as described previously (9). Briefly, the x-axis of the grid was centered on the soma and the y-axis was aligned with the second row of the grid placed on the L1/2 border.

About 10% of all patched neurons across all layers were presumptive inhibitory cells as evidenced by membrane capacitance < 100 pF and high levels of spontaneous synaptic input. These were excluded from the data set. Cells with resting Vm > 50 mV, those that exhibited spontaneous repetitive firing, or whose passive membrane properties fluctuated significantly over the course of data collection were also excluded.

#### Excitation profiles

We measured the excitability across layers (i.e. number of APs per UV flash) and how far from the soma a UV flash can evoke an action potential (AP), using cell-attached recordings to detect APs. We sampled cells from all cortical layers to measure excitation profiles (n = 20 for each hemisphere). To map excitation profiles, we used an 8×8 grid with 50 μm spacing for L2, L3, L4 and L6 neurons and for L5 pyramidal neurons we used an 8×16 grid with the same spacing to test for dendritically evoked spiking. Action potentials within 50 ms of the UV flash onset were included in the analysis. Using these excitation profiles we determined that a laser power between 30-35 mW evoked reliable synaptic responses in all neurons recorded. The same laser power was used for synaptic maps and excitation profiles (30-35 mW).

### Analysis of LSPS data

Analysis was conducted as described previously (9). Briefly, the mean current amplitude of synaptic events were calculated in the 50ms epoch after the direct response time window (7.5 ms after UV stimulus). Direct responses triggered by stimulation around somata were excluded in our analyses. We recorded 2 to 4 maps for each cell to create an average input map, and these average maps were used for group averages and for all analyses. Like the individual maps, population maps were aligned with respect to the soma on the x axis and the L1/L2 boundary on the y axis. Pixels in the population maps where direct responses were recorded in >50% of cells, i.e. areas proximal to the cell body, were excluded from the analysis.

To create input-output functions from the map data, average maps for each cell were summed over either the antero-posterior axis or the laminar axis to yield 1×16 vectors of input strength vs. location. These vectors were then binned according to laminar or antero-posterior location, respectively, and combined to yield n x 16 or 16 x n matrices relating presynaptic input to postsynaptic soma location (“output”), where n = number of bins as reported in the individual figure legends. The matrices were 2D-interpolated for display purposes.

*Statistical significance between hemispheres* for population map values was computed pixel-by-pixel, with the average map value for each cell being considered a single data point. Comparisons were made via 2-tailed, unpaired t-tests. For comparison, randomized datasets were generated by pooling all maps from both hemispheres for each layer and drawing maps at random to give two pools with the same n as the original data sets.

### In vivo recordings

#### Subjects and Surgery

14 CBA/J mice, aged P30-60, were used in accordance with the National Institute of Health guidelines, as approved by the City College of New York Institutional Animal Care and Use Committee. We administered ketamine (75mg kg^-1^)/medetomidine (0.5mg kg^-1^) before a stereotactic surgery was performed to allow for awake recordings. The scalp from the entire top of the skull was removed to reveal Bregma and Lambda, barely exposing the muscle of the temporal bone. A metal plate was bound to the exposed bone via two layers of Metabond and a single layer of Vitribond, followed by a covering of dental cement. Following administration of ketamine (75mg kg^-1^)/medetomidine (0.5mg kg^-1^) mice were head-fixed on a freely rotating wheel inside a sound-proof chamber. We made a small craniotomy and durotomy over the auditory cortex. The position of the auditory cortex was determined by the relative positions of bregma and lambda, as well as the specific bone sutures of the temporal bone. Craniotomies were 0.1-0.2mm in diameter to provide stability to our recordings. Mice were allowed at least 24 hours post-surgery to recover before any recording sessions began.

#### Electrophysiology

We recorded from the Left and Right ACx of awake mice. During the recording sessions we targeted neurons in superficial layers (L2/3, L4; 150-450 microns below cortical surface) using the standard blind cell-attached technique (39). Electrodes were pulled from a glass borosilicate filament and filled with either physiological saline (0.9% NaCl) or intracellular solution ((in mM) 128 K-methylsulfate, 4 MgCl, 10HEPES, 1EGTA, 4NaATP, 0.3NaGTP, 10 Na-phos-phocreatine) and had resistances between 4-8MegaOhms. Recordings were obtained using Axopatch 200B (Axon Instruments) and custom electrophysiological data acquisition software (exper, Hromadka) written in MATLAB (Mathworks). We detected the presence of cells in the cortex based on changes in pipette resistance.

A soundproof chamber was used to conduct all recordings. We used a custom built real-time Linux system (200 kHz sampling rate) driving a high-end Lynx L22 audio card (Lynx Studio Technology Inc., Newport Beach, CA) with an ED1 electrostatic speaker (Tucker-Davis Technologies, Alachua, FL) in free-field configuration (speaker located 6 inches lateral to, and facing, the contralateral ear). The stimuli were generated with custom MATLAB scripts. To compute tuning curves of our cells’ best frequency we used a set of pure tones that lasted 100ms long of 16 different frequencies at 3 intensity levels (20dB, 50dB, 80dB). Tuning curves were only used to test the position of our recordings to ensure that we were within the ACx.

### Time constant posterior distributions

#### Autocorrelation and single neuron time constant estimation

For this analysis, we used recordings in which no auditory stimulus was played. We binned the recordings in Δ=20ms bins and computed the autocorrelation of the binned signal. For each neuron, we performed least squares fits of exponential decays of the form

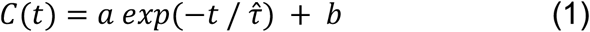

with multiple initializations to extract the values of the parameters 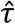 and a. We assumed the process is stationary and fixed the value of b to (λΔ)^2^ where λ is the mean firing rate. We then estimated the bias and variance of the time constant 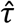 for each neuron using the Dichotomized Gaussian model.

#### Dichotomized Gaussian model

The Dichotomized Gaussian (DG) model is a spiking generative model that can capture arbitrary correlation coefficients in spiking neural data (40). Sample spike trains X(t) are generated by thresholding samples of a latent multivariate Gaussian time series V(t)

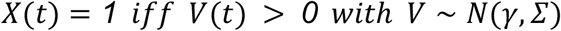

with Σ(t, t)=1 without loss of generality. Assuming the process is stationary, γ is a constant and the time covariance matrix Σ(t, s)=Σ(t - s) is only a function of the lag between time points. The latent parameters γ and Σ are obtained from the spiking rate λ and autocorrelation C using

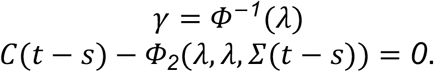

The equation for Σ(t - s) is monotonic on the variable and has a unique solution constrained to be in the range (−1, 1). In this way, the DG model can capture specific spiking rates and temporal auto-correlations.

#### Bias and variance estimation

For each neuron, we generated 400 sets of surrogate data replicating the recording conditions and using the DG model with the exponential autocorrelation of equation (1) and the extracted parameters 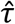 and A. We repeated the estimation procedure for each set and obtained a set of surrogate time constants {τ^DG^_1_,τ^DG^_2_, …,τ^DG^_400_}. We assumed these values are observations from a log-normal distribution 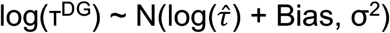 and we estimate its mean and variance by maximum likelihood. The difference between the estimated mean E[log(τ^DG^)] and 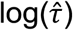 is an estimate of the bias of our estimator in our procedure and σ^2^ represents the uncertainty of the estimate.

#### Evidence integration

We assumed that the bias corrected time constants 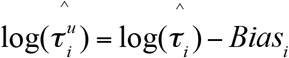 obtained from the neurons in each hemisphere come from a log-normal distribution 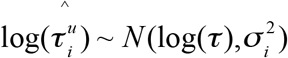 with *τ* the network’s time constant and 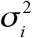 the variance estimated for each neuron. Assuming a uniform prior for *τ*, we integrate the evidence from the observations of each hemisphere into the posteriors by computing

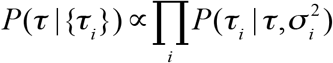

## References

1. Albouy P, Benjamin L, Morillon B, Zatorre RJ. Distinct sensitivity to spectrotemporal modulation supports brain asymmetry for speech and melody. Science. 2020;367(6481):1043–7.

2. Boatman DF, Miglioretti DL. Cortical sites critical for speech discrimination in normal and impaired listeners. The Journal of neuroscience : the official journal of the Society for Neuroscience. 2005;25(23):5475–80.

3. Boatman D. Cortical bases of speech perception: evidence from functional lesion studies. Cognition. 2004;92(1-2):47–65.

4. Arnal LH, Poeppel D, Giraud AL. Temporal coding in the auditory cortex. Handb Clin Neurol. 2015;129:85–98.

5. Giraud AL, Poeppel D. Cortical oscillations and speech processing: emerging computational principles and operations. Nat Neurosci. 2012;15(4):511–7.

6. Levy RB, Marquarding T, Reid AP, Pun CM, Renier N, Oviedo HV. Circuit asymmetries underlie functional lateralization in the mouse auditory cortex. Nat Commun. 2019;10(1):2783.

7. Wetzel W, Ohl FW, Scheich H. Global versus local processing of frequency-modulated tones in gerbils: an animal model of lateralized auditory cortex functions. Proc Natl Acad Sci U S A. 2008;105(18):6753–8.

8. Shepherd GM, Svoboda K. Laminar and columnar organization of ascending excitatory projections to layer 2/3 pyramidal neurons in rat barrel cortex. The Journal of neuroscience : the official journal of the Society for Neuroscience. 2005;25(24):5670–9.

9. Oviedo HV, Bureau I, Svoboda K, Zador AM. The functional asymmetry of auditory cortex is reflected in the organization of local cortical circuits. Nat Neurosci. 2010;13(11):1413–20.

10. Goldman MS. Memory without feedback in a neural network. Neuron. 2009;61(4):621–34.

11. Hooks BM, Hires SA, Zhang YX, Huber D, Petreanu L, Svoboda K, et al. Laminar analysis of excitatory local circuits in vibrissal motor and sensory cortical areas. PLoS Biol. 2011;9(1):e1000572.

12. Weiler N, Wood L, Yu J, Solla SA, Shepherd GM. Top-down laminar organization of the excitatory network in motor cortex. Nat Neurosci. 2008;11(3):360–6.

13. Gaese BH, Ostwald J. Complexity and temporal dynamics of frequency coding in the awake rat auditory cortex. Eur J Neurosci. 2003;18(9):2638–52.

14. Wilting J, Priesemann V. Inferring collective dynamical states from widely unobserved systems. Nat Commun. 2018;9(1):2325.

15. Zeraati R, Engel, T. A., & Levina, A. Estimation of autocorrelation timescales with Approximate Bayesian Computations. BioRxiv. 2020.

16. Marlin BJ, Mitre M, D’Amour J A, Chao MV, Froemke RC. Oxytocin enables maternal behaviour by balancing cortical inhibition. Nature. 2015;520(7548):499–504.

17. Ehret G. Left hemisphere advantage in the mouse brain for recognizing ultrasonic communication calls. Nature. 1987;325(6101):249–51.

18. Zylberberg J, Strowbridge BW. Mechanisms of Persistent Activity in Cortical Circuits: Possible Neural Substrates for Working Memory. Annu Rev Neurosci. 2017;40:603–27.

19. Seung HS, Lee DD, Reis BY, Tank DW. Stability of the memory of eye position in a recurrent network of conductance-based model neurons. Neuron. 2000;26(1):259–71.

20. Hart E, Huk AC. Recurrent circuit dynamics underlie persistent activity in the macaque frontoparietal network. Elife. 2020;9.

21. Chaudhuri R, Knoblauch K, Gariel MA, Kennedy H, Wang XJ. A Large-Scale Circuit Mechanism for Hierarchical Dynamical Processing in the Primate Cortex. Neuron. 2015;88(2):419–31.

22. Schuecker JG, S., Helias, M.. Optimal Sequence Memory in Driven Random Networks. Physical Review X. 2018;8(4), 041029.

23. Kennedy A, Kunwar PS, Li LY, Stagkourakis S, Wagenaar DA, Anderson DJ. Stimulus-specific hypothalamic encoding of a persistent defensive state. Nature. 2020;586(7831):730–4.

24. Murray JD, Bernacchia A, Freedman DJ, Romo R, Wallis JD, Cai X, et al. A hierarchy of intrinsic timescales across primate cortex. Nat Neurosci. 2014;17(12):1661–3.

25. Wasmuht DF, Spaak E, Buschman TJ, Miller EK, Stokes MG. Intrinsic neuronal dynamics predict distinct functional roles during working memory. Nat Commun. 2018;9(1):3499.

26. Cavanagh SE, Towers JP, Wallis JD, Hunt LT, Kennerley SW. Reconciling persistent and dynamic hypotheses of working memory coding in prefrontal cortex. Nat Commun. 2018;9(1):3498.

27. Cavanagh SE, Wallis JD, Kennerley SW, Hunt LT. Autocorrelation structure at rest predicts value correlates of single neurons during reward-guided choice. Elife. 2016;5.

28. Spitmaan M, Seo H, Lee D, Soltani A. Multiple timescales of neural dynamics and integration of task-relevant signals across cortex. Proc Natl Acad Sci U S A. 2020;117(36):22522–31.

29. Fiser J, Chiu C, Weliky M. Small modulation of ongoing cortical dynamics by sensory input during natural vision. Nature. 2004;431(7008):573–8.

30. Hung CP, Ramsden, B. M., & Roe, A. W.. Inherent biases in spontaneous cortical dynamics. The Dynamic Brain: An Exploration of Neuronal Variability and Its Functional Significance: Oxford University Press; 2010. p. 83–103.

31. Poeppel D. The analysis of speech in different temporal integration windows: Cerebral lateralization as ‘asymmetric sampling in time.’. Speech Communication. 2003;41(1):245–55.

32. Doupe AJ, Kuhl PK. Birdsong and human speech: common themes and mechanisms. Annu Rev Neurosci. 1999;22:567–631.

33. Shepard KN, Lin FG, Zhao CL, Chong KK, Liu RC. Behavioral relevance helps untangle natural vocal categories in a specific subset of core auditory cortical pyramidal neurons. The Journal of neuroscience : the official journal of the Society for Neuroscience. 2015;35(6):2636–45.

34. Hertz S, Weiner B, Perets N, London M. Temporal structure of mouse courtship vocalizations facilitates syllable labeling. Commun Biol. 2020;3(1):333.

35. Marconi MAN, D; Abbasi, R.; Penn, D.J.; Zala, S.M.. Ultrasonic courtship vocalizations of male house mice contain distinct individual signatures. Animal Behaviour. 2020;169:169–97.

36. Oviedo HV. Connectivity motifs of inhibitory neurons in the mouse Auditory Cortex. Sci Rep. 2017;7(1):16987.

37. Neophytou D, Oviedo HV. Using Neural Circuit Interrogation in Rodents to Unravel Human Speech Decoding. Front Neural Circuits. 2020;14:2.

38. Suter BA, O’Connor T, Iyer V, Petreanu LT, Hooks BM, Kiritani T, et al. Ephus: multipurpose data acquisition software for neuroscience experiments. Front Neural Circuits. 2010;4:100.

39. Hromadka T, Deweese MR, Zador AM. Sparse representation of sounds in the unanesthetized auditory cortex. PLoS Biol. 2008;6(1):e16.

40. Macke JH, Berens P, Ecker AS, Tolias AS, Bethge M. Generating spike trains with specified correlation coefficients. Neural Comput. 2009;21(2):397–423.

